# High-throughput Image-based Clustering of CAR-T/Tumor Cocultures for Rapid and Facile Hit Identification

**DOI:** 10.1101/2024.08.28.609577

**Authors:** Zhi Xu, Xueqi Liu, Kirby Madden-Hennessey, Jordan Urbani, Shahryar Khoshtinat Nikkhoi, Anusuya Ramasubramanian, Kartika G. Venugopal, Qi Zhao, Eric L. Smith, Yun Fu

## Abstract

Chimeric antigen receptor T cell is important because of its potential to treat various diseases. As deep learning continues to advance, using unsupervised methods to classify medical images has become a significant focus because collecting high-quality labeled data for medical images is labor-intensive and time-consuming. Beyond the need for accurate labeling, there is a desire to explore the underlying characteristics of the data, even when labels may be ambiguous or uncertain. To address these challenges, we present a novel approach that combines image clustering with an insightful explanation of how these clusters are formed. Our method employs a U-net combined with a clustering algorithm to segment the dataset into different groups. After clustering, we use various techniques to interpret and elucidate the results. Moreover, our paper introduces a unique dataset focused on cell data, specifically highlighting the developmental patterns of cancer cells and T cells under various experimental conditions. This dataset offers a rich source of information and presents a complex challenge for image classification due to the diversity of conditions and cell behaviors involved. Our study thoroughly compares different architectural models on this new dataset, demonstrating the superior performance of our proposed architecture. Through experimental analysis and ablation studies, we provide substantial evidence of the benefits offered by our architecture, not only in terms of accuracy but also in its ability to reveal deeper insights into the data. This work advances the field of image classification and opens new possibilities for understanding complex biological processes through computer vision.

## 1 Introduction

Chimeric antigen receptor T (CAR T) cell therapy has become a significant focus in immunotherapy research because of its potential to treat various diseases. One of its crucial aspects is the integration of several components, including the receptor, spacer/transmembrane, and intracellular signaling domains. Interestingly, the binding affinity of the domain, such as the scFv or VH domain, doesn’t always correlate with the CAR T’s effectiveness [1], and minor modifications in the design can lead to unexpected changes in the toxicity and behavior of CAR T cells [2]. Identifying which changes within the construct produce persistent CAR T cells is important in determining the CAR T construct that is suitable for clinical translation [3]. It has been noticed that patients who undergo a relapse following CAR T cell therapy often show a lack of long-term persistence of their CAR T cells. [3]. However, predicting the CAR T construct that shows the greatest persistence *in vivo* has proven challenging. Current assays for identifying highly persistent CAR T candidates often suffer from two major pitfalls. These assays that rely on a single or few stimulations with antigen-presenting target cells are time-saving but unable to accurately identify highly persistent phenotypes. Assays require tedious reestablishment of tumor and CAR T cell cocultures are accurate but costly and long to execute. In addition, many of these assays are limited in throughput, reducing the number of candidates that can be effectively screened[4]. This often restricts evaluations to fewer candidates [5] [6]. Given the limitations of current methods, this study introduces the Trovo system, a new coculture imaging workflow, designed to work alongside deep learning, that allows for the semi-pooled screening of large libraries of CAR T candidates. Deep learning offers a way to process large data sets and could help understand the nuances of T cell/tumor co-culture interactions.

Despite the notable advancements achieved thus far in processing large data sets from high-throughput screens, drug developers have expressed the need for comprehensible text explanations accompanying the outcomes of deep learning models. The integration of interpretability serves a dual purpose: enhancing the grasp of deep learning results and bolstering the models’ dependability. With explanatory text, researchers can readily identify any weaknesses or limitations within the model’s results, thereby harnessing the model’s capabilities more effectively in their development process. As illustrated in Fig. 1, our proposed cluster methods not only assign images to distinct clusters but also provide visual and textual explanations for each cluster.

**Figure 1:**
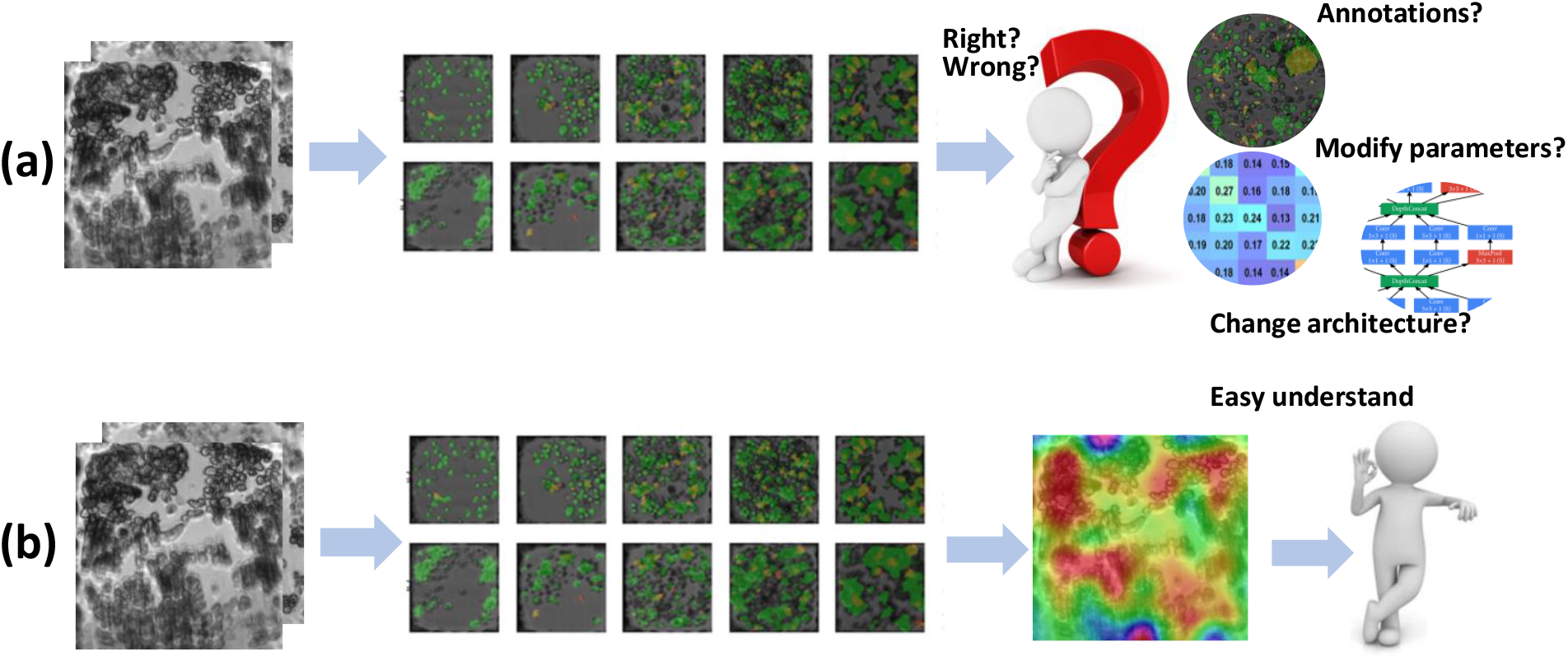
**A**: Given input images, we can use a clustering algorithm to label each image into a cluster. However, without ground truth, the correctness of the results is unconvincing, and it is not easy to improve the algorithm. **B**: With explanations, the result of the algorithm is more convincing and easy to understand.

Specifically, using Grad-CAM, we can identify the regions of interest in neural networks without retraining. Additionally, the emergence of large language models (LLMs) allows us to generate templates and input information to LLMs to analyze the influence of different environments. By integrating these technologies, our model can provide both visual and textual explanations, leveraging the data collected from Trovo enables us to expand our capabilities, aiding experts in understanding the effects of different variables.

To achieve image classification and provide explanations, our model comprises several components: a feature extractor, a clustering algorithm, and an explanation module. Through rigorous experimentation, we confirm the effectiveness of our proposed architecture and the accompanying explanation module. Furthermore, we present intra- and inter-covariance matrices as supplementary evidence to support the efficacy of our clustering algorithm. Visualizations of the results demonstrate the proficiency of our explanation modules in a graphical manner.

In brief, our contributions include:

- The introduction of a novel dataset, coupled with the proposal of an architectural framework adept at extracting vital information from input microwell images. A multi-head attention layer is strategically incorporated to synergize U-net-extracted and human-designed features.
- The formulation of two distinct explantion modules, crucial in facilitating a deeper understanding of our model’s cluster results. The visual explanation module serves as a guide to discern where the model directs its focus. In contrast, the text explanation module highlights the unique characteristics of clusters and distinguishes them from their counterparts.
- The execution of an extensive array of experiments, designed to showcase the efficiency of our proposed explanation modules. The validation process extends to incorporating intra and inter-covariance matrices, comprehensively evaluating the cluster algorithm’s effectiveness.

## 2 Related Works

### 2.1 Medical Image Classification

Since the development of deep learning, more and more researchers are trying to use deep learning-based methods to help them solve medical problems, including medical image classification. U-Net ([7]) proposed by Olaf, which is based on FCN ([8]), is a widely-used architecture in medical imaging. The use of the skip connection makes full use of the features of the encoder. Recently, several experts within the field have focused on using self-supervised or semi-supervised methods to advance the effectiveness of medical image classification. For example, Azizi et al. [9] noticed that a big self-supervised model can greatly enhance the effectiveness of medical image classification. SRC [10] introduces sample relation consistency to help better extract information from the unlabeled medical data. Lei Zhou et al. [11] tries to use self pre-training based on Mask Autoencoder ([12]) to improve the effectiveness of medical image classification. In this work, we tested several different architecture which can combine extracted features and human-designed features to improve the accuracy of the model.

### 2.2 Image clustering

Image clustering is an unsupervised machine-learning technique focused on finding similar features between different samples and labeling each sample into a cluster. The most typical image clustering algorithm is K-means ([13]), which was proposed in 1956. Recently, with computer vision and data mining development, several clustering algorithms have been proposed. For example, Yunfan Li et al. [14] tries to use Contrastive learning to perform image clustering and achieve reliable outcomes on multiple datasets. In another example, Shakeel et al. [15] introduces a method to improve the performance of clusters on lung cancer detection. This paper uses an encoder and a multi-head attention module to better extract image features and improve the clustering results.

### 2.3 Explainable deep learning

As deep learning continues to advance, researchers are only satisfied with using neural networks if they understand why a decision is made. Hence, some researchers focus on the multiple ways of explaining the output of neural networks. One way is to generate an attention map based on the input image. Bolei Zhou et al. [16] initially did this with Class activation mapping, and since then, researchers have generated better attention maps like Grad-CAM ([17]), ScoreCAM ([18]), RelevanceCAM ([19]), PullzeCAM ([20]). Another way to explain the output of the neural network is the concept-based method, which was introduced by TCAV ([21]). Ghorbani et al. [22] tries to automatically find the explanation based on the concept. Ge et al. [23] build a graph neural network (GNN) to better understand the network’s decision reasoning. In this paper, we provide visual and text information to explain why the network makes a particular decision and help researchers understand where the network is focused.

## 3 Methods

Our proposed methods consists of four different parts: a feature extractor to extract the features for every group of images, a cluster module to label each group of images to a cluster, a visual explanation module that uses Grad-Cam to provide a visual explanation, and a text explanation module which uses the features and templates made by the large language model to generate text explanation. Fig.2 demonstrates the whole architecture of the proposed methods.

**Figure 2:**
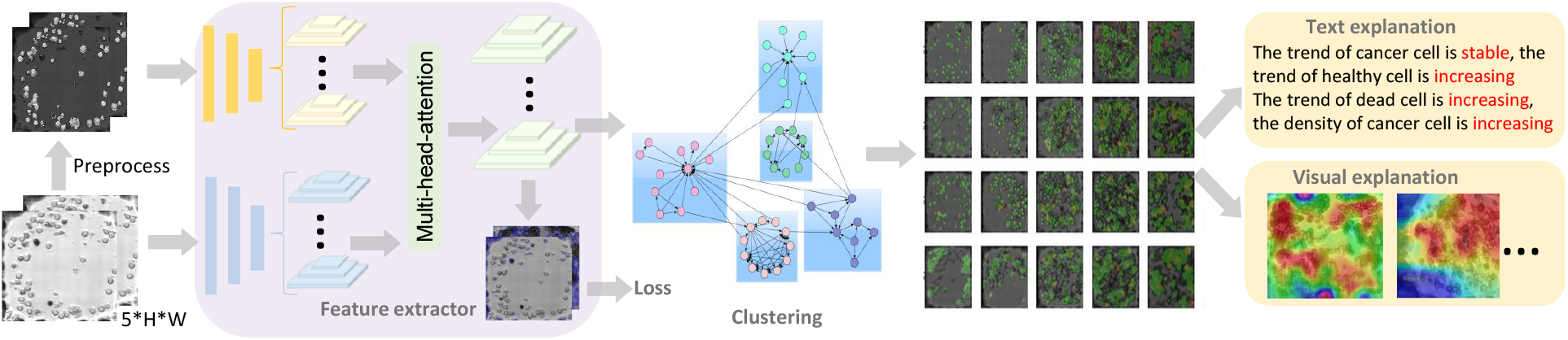
Given input images, our model will first use a feature extractor to aggregate information from the input images; then we will use a clustering algorithm to label each image into a cluster. After getting multiple clusters, we will use text explanation and visual explanation methods to explain why we cluster each image.

### 3.1 Feature extractor

Our advanced feature extraction framework is a multi-faceted system, intricately engineered to conduct a detailed analysis of cellular images. This framework is composed of several critical elements, each playing a significant role in processing and understanding the complex data presented in these images.

Central to our framework is a CNN encoder, which is exquisitely tailored to delve into and extract the complex, nuanced information hidden within each image. This encoder is not working in isolation; it is complemented by a multi-head attention module. This module is adeptly integrated into our system to uncover and interpret the temporal nuances and patterns evident across sequences of images.

To further enhance our system’s capability, we have incorporated a custom-designed feature extractor. This extractor is specifically developed to identify and isolate crucial low-level features from each image. By doing so, it provides a comprehensive and detailed representation of the data, capturing essential aspects that might otherwise go unnoticed.

As shown in Fig 2, we show a detailed visual depiction of the architecture to provide a more in-depth understanding of our feature extraction framework. A key aspect of our feature extraction process is the utilization of both provided annotations and the integration of pseudo-labels. The pseudo-labels are derived from the analysis of T cells and cancer cells within the images, offering a novel means of enhancing our system’s effectiveness, even without the labeled data.

The creation of these pseudo-labels is based on an essential observation: the cellular regions in the images exhibit unique brightness characteristics, distinguishing them from the background. By converting images into the HSV color space, we can precisely separate these regions, enabling the accurate generation of pseudo-labels and thereby enriching the quality of our input data.

In addition, our framework employs a variety of human-designed features, such as the density and proliferation rate of cancer and T cells. These features provide valuable statistical insights, which are often challenging for neural networks to capture autonomously.

The culmination of our feature extraction process is marked by the integration of the multi-head attention module. This module is specifically designed to combine insights gleaned from different images. It excels in capturing the evolving temporal trends within groups of images and effectively combining them with human-designed features.

By integrating all these components, our model is adept at extracting a multitude of information layers from the original input, thereby generating a rich and informative feature set for subsequent clustering. This comprehensive approach ensures our model’s efficacy in not just classifying but also understanding the patterns present in cellular imaging data.

### 3.2 Cluster module

Upon successful training of the feature extractor, the remaining challenge is the assignment of labels to groups of images, thereby organizing them into coherent clusters. However, a notable hurdle arises from the absence of labels regarding which image groups should belong to the same cluster. This complication precludes the direct training of a model for group clustering. Thus, a search for a suitable clustering method ensues, one that can adeptly assign inputs to multiple clusters while the cluster count is unknown.

We selected the AffinityPropagation algorithm ([24]) as our clustering approach. A few advantages of AffinityPropagation include its ability to circumvent the requirement for a predetermined cluster count and its resilience to the influence of initial values. This is a distinct contrast to algorithms like K-means. However, a significant consideration emerges due to the nature of our feature extractor’s output – a high-dimensional feature space rife with dimensions that may prove extraneous for clustering purposes. To address this, a preliminary step involves dimensionality reduction to eliminate redundant dimensions. To this end, we employ T-SNE (t-Distributed Stochastic Neighbor Embedding) ([25]), a method that facilitates the transformation of complex high-dimensional data into lower-dimensional data that is conducive to subsequent clustering.

Having reduced the dimensionality of the data, the application of the AffinityPropagation algorithm becomes more feasible. The resultant application of AffinityPropagation yields multiple clusters, each containing a grouping of image sets. However, because of the absence of the labeled data, assessing the performance and quality of these clusters is not a straightforward task.

To surmount this challenge and provide insights into the efficacy of the cluster results, two distinct explanation modules were developed. These modules are specifically designed to elucidate the clustering outcomes’ rationale by enhancing the cluster assignments’ interpretability. To do this, we present visual and text explanation modules. The former serves as a mechanism to highlight the key focus areas within each cluster and employs attention maps to emphasize regions of significance. Meanwhile, the text explanation module articulates each cluster’s defining characteristics and elucidates the distinctions between clusters. By seamlessly integrating these two explanation modules, we augment the understanding of our clustering methodology and address the challenge posed by the absence of ground truth.

### 3.3 Visual Explanation Module

In this section, we delve into the methodology behind our generation of visual explanations for each cluster. The objective of our visual explanation module is to discern the focal points of our model’s attention within each image, subsequently aiding user understanding of the appropriateness of our model’s cluster assignments based on the input images. Taking inspiration from Grad-Cam ([17]), we leverage gradient information to create attention maps.

However, a noteworthy challenge emerges: our model lacks individual labels for each image, which complicates the direct utilization of gradients derived from target concepts. To circumvent this issue, we adopt a strategic approach post-clustering. Specifically, we assign a cluster index to each image, effectively creating pseudo-labels. Subsequently, we retrain a classifier solely based on the extracted features. Notably, during this classifier training phase, we exclusively update the classifier while keeping the feature extractor unchanged. This deliberate decision maintains the feature extractor’s consistency and integrity, safeguarding it against potential distortion by potentially erroneous pseudo-labels.

Following the training of the classifier, we emulate the Grad-Cam methodology by employing a similar configuration. In this context, the target cluster index is employed as the target concept, and the attention map is computed based on the original images. Mathematically, this process can be formalized as follows:

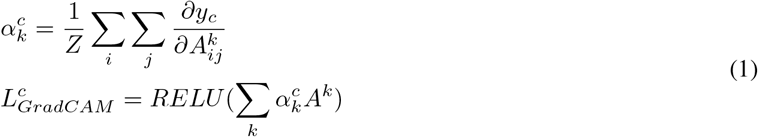

Here, 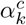 represents the importance weight attributed to each feature map *A*_*k*_. The significance of these feature maps is aggregated through summation, followed by filtration of extraneous information using a rectified linear unit (*RELU*) function. This process culminates in the creation of an attention map that highlights the pertinent regions within the original images, effectively elucidating the visual cues that our model utilizes for its cluster assignments.

### 3.4 Text Explanation Module

Within this module, our focus shifts towards delineating the process by which we generate text explanations tailored to each cluster. The underlying goal is to provide user-readable explanations that facilitate a clear understanding of the rationale behind our model’s cluster assignments.

To achieve this, we constructed a template that serves as a structured framework for generating text explanations. This template contains multiple dimensions of information, encompassing key aspects such as cell densities, proliferation trends, and comparative analysis with other clusters. With a template in place, the subsequent challenge involves populating it with relevant information.

In an effort to bolster the robustness of our explanatory model, we take a novel approach: sidestepping the direct utilization of cancer and T Cell annotations. Instead, we employ these annotations to train an auxiliary model that predicts cell density based on input images. Leveraging the predictions from this auxiliary model, we gain the ability to generate explanations even in the absence of explicit annotations. This strategic move enhances our approach’s adaptability and contributes to the overall resilience of our model’s performance.

Similar to the approach taken for generating density-based explanations, the generation of explanations pertaining to proliferation trends follows a comparable strategy. Here, a model is trained to predict proliferation trends, thus enabling the prediction of related information for explanation.

A distinguishing feature of our text explanations lies in their comparative analysis. We adopt a comparative perspective to highlight each cluster’s unique attributes. Specifically, we assess the cluster that exhibits the most distinct characteristics compared to other clusters. This distinctiveness forms the explanatory text’s crux, emphasizing the distinguishing traits that underlie each cluster’s composition.

The resultant text explanations, enriched by these comprehensive elements, allow users to discern the disparities and nuances between clusters readily. Through this mechanism, our methodology transcends the realm of statistical clustering, fostering an in-depth understanding of the intrinsic features that set each cluster apart.

## 4 Experiments

### 4.1 Experiments setup

#### 4.1.1 Trovo system

The Trovo system leverages the benefits of a conventional petri dish or microtiter plate bulk cultures and transforms these platforms into thousands of microchambers, enabling easy operation, compatibility with existing protocols, and optimal cell health. The hydrogel-based well system segregates cells while maintaining rigidity, transparency, and customizability. Our microwell generation software is fully parameterized, producing 300-micron-high walls and 2.5 k microwells in 10 minutes. The Trovo, which can be seen in Fig.3, uses hydrogel lithography, allowing customizable microwell size and shape. Essentially, a 405 nm laser induces the gelation of various photopolymerizable and enzymatically degradable hydrogels. The optimal compositon of the hydrogel forms optically transparent grids that bind well to glass bottom culture dishes. For suspension cultures, cells at the bottom of microwells are unaffected by macroscopic disturbances such as medium changes. This micro-trapping property ensures high-density clone segregation and the registration of clone behavior.

**Figure 3:**
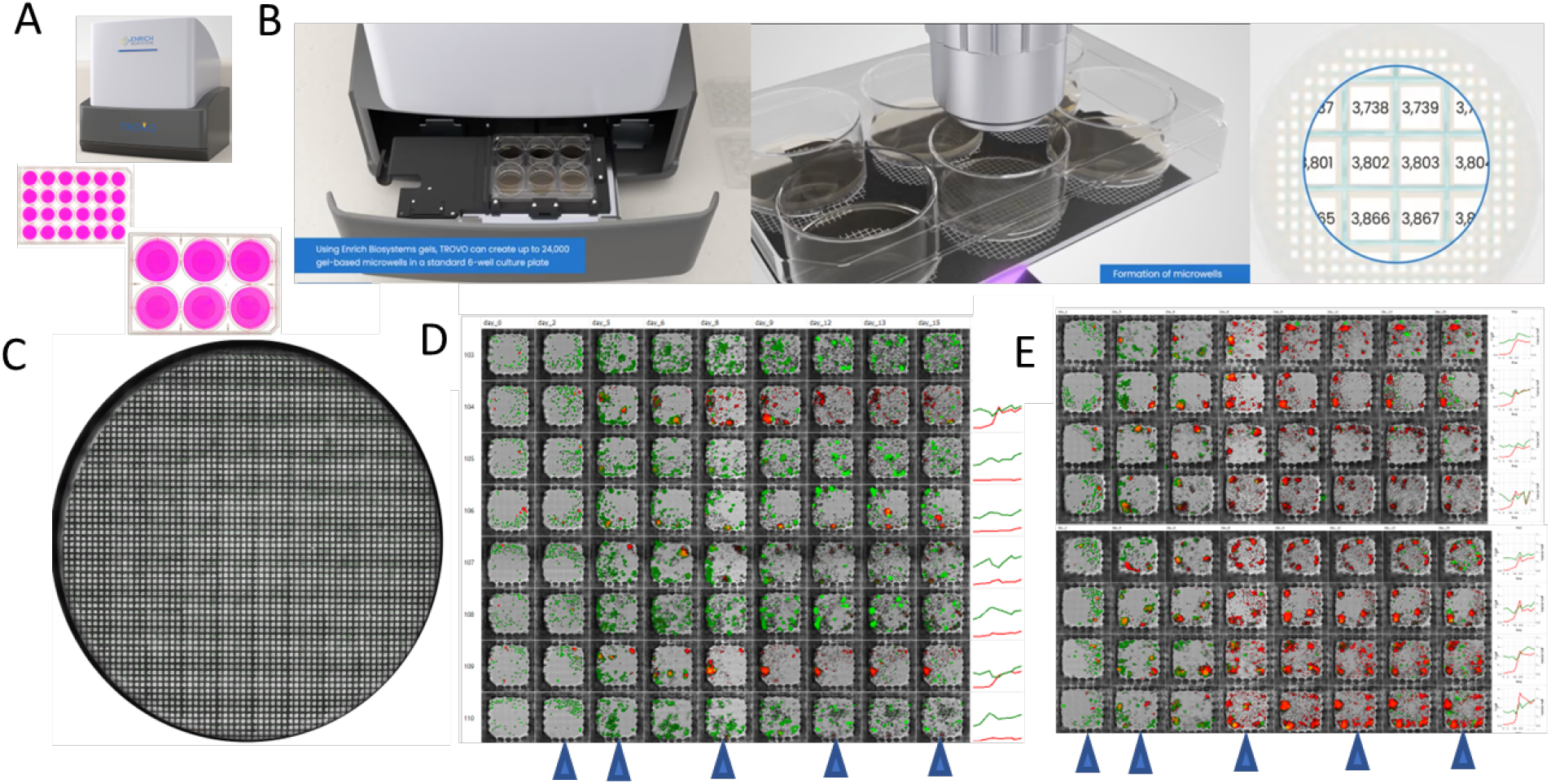
Screening long-term tumor killing and T cell proliferation/persistence behavior of CAR T clones using Trovo. (A) The hydrogel microwell system can be made on a 6-well or 24-well culture dish using the Trovo system. (B) T-cell tumor co-cultures are seeded into each microwell, observed daily, analyzed, and isolated based on the behavior of each microwell. (C) A “bird’s-eye” view of a typical microcompartment co-culture on day 8. Each microwell is a 250 *µ*m x 250 *µ*m square. (D, E) Collection of microwell images over several weeks showing distinct clonal behavior. The time series data of microchambers were typically collected from 2300 chamber experiments on Trovo. (Yellow/Red: T cells, Green: live tumor, Grey: dead tumor).

#### 4.1.2 Microwell Printing

About 2500 microwells were printed within each well’s central 20mm glass bottom area within a 6 well glass-bottom microplate. Microwell printing gel (Enrich Biosystems) was added to each well and placed in the Trovo instrument (Enrich Biosystems) for printing. After a 1 hr automated printing process, 2 mL of 1X PBS was added to the wells and incubated at 37°C for 5 minutes to wash. This washing step was repeated three times prior to the addition of complete medium for overnight incubation at 37°C.

#### 4.1.3 CAR Generation Methodology

To establish retroviral producer lines, H29D cells underwent transfection with retroviral vectors encoding three BCMA-targeting CARs, CAR C (BCMA-CD28-CD3z), CAR A (BCMA-41BB-CD3z), CAR B (BCMA-CD3z), Del (BCMA scFv-deleted signaling domains) utilizing TransIT-293 (Mirus Bio). Following a 72-hour incubation period, the resulting supernatant was harvested, subjected to filtration, and used to infect GalV9 cells. After 5 days, 600,000 GalV9 producer cells were plated in a 6-well plate. Supernatant containing the retrovirus was collected at 24 and 48-hour intervals. For transduction, T cells were activated utilizing TransAct (Miltenyi Biotec), CD3 and CD28 agonists, along with recombinant human IL-2 (rhIL-2; 100 U ml-1; Proleukin), IL7 (60 ng ml-1; PeproTech), and IL15 (10 ng ml-1; PeproTech), in complete media (RPMI 1640, 10% FBS, 1% Gluramax, and 1% Penicillin-Streptomycin) overnight. Subsequently, activated T cells were plated at a density of 500,000 cells per well on Retronectin (Takara)-coated 24-well plates and supplemented with twice the concentration of cytokines. Following the addition of viral supernatant, cells were subjected to centrifugation (2000xg, 1 hour) and incubated at 37°C in a 5% CO_2_ environment. CAR expression on T cells was accessed by flow cytometry 7 days post activation by sequentially staining with a flag-tagged BCMA protein (made in-house) and an anti-FLAG tag antibody conjugated with PE-Cyanine 7 (Biolegend Cat. 637323).

#### 4.1.4 Co-Culture Setup

OPM2, a myeloma cell line, labeled with GFP (OPM2-GFP), and three different CAR T cells constructs stained with CD4 Alexa fluor 647 (1:200, Biolegend, cat. #344636) and CD8 Alexa fluor 647 (1:200, Biolegend, cat. #301022) were combined and pooled at a final E:T ratio of 1:10 in a final volume of 200 uL. Phenol Red-free Medium (Thermo Fisher, cat. #11835030), 20% FBS (Thermo Fisher, cat. #10082147), 100 U/mL Pen/Strep (Thermo Fisher, cat. #15140122), 1X Glutamax (Thermo Fisher, cat. #35050061) were removed from the wells, and the cell mixture was added drop wise throughout the respective 20 mm microwell. The plate was incubated for 15 minutes at 37°C to allow the cells to settle. Once the incubation was completed, 1.5 mL of complete medium was added to the dish and incubated at 37°C for 1 hr. Following the incubation, the plate was imaged using the Trovo system. On subsequent days, CAR T cells were stained by removing all media and adding a CD4/CD8 mastermix drop wise to the center of the well(s) prior to imaging. This was repeated on the following days: 0, 1, 2, 3, 6, and 7. A chronic antigen exposure assay was performed on day 3 of the experiment. After imaging, the medium was removed and 25K OPM2-GFP cells were added dropwise to the 20 mm glass bottom area. Cells were incubated for 15 minutes to allow the cells to settle before adding 1.5 mL of complete medium.

The above experiment was repeated for each coculture containing a CAR T cell candidate and OPM2-GFP target cells. CAR T cells were stained for 1 hr with CD4/CD8 Alexa Fluor 647 prior to imaging on the following days: 0, 2, 3, 6, 7, and 8. Similar to above, a chronic antigen exposure test was performed on days 3 and 6. After imaging on their respective days, the medium was removed and 25K OPM2-GFP cells were added dropwise to the 20 mm glass bottom area. After adding 1.5 mL of complete medium, the cells were incubated for 15 min to allow the cells to settle. The microwell imaging dataset was analyzed as described below.

### 4.2 Data Description

In this article, we introduce a novel dataset that delves into the specific characteristics that distinguish our data set from others in the field. Fig 5 provides a visual representation of the dataset, showcasing the developmental trajectories of cancer cells and T cells.

Our dataset is organized into numerous subsets, each comprising a series of five distinct images. These images represent microwells captured over consecutive days, offering a dynamic, temporal view of cellular behavior. For each image, we have annotated two key types of cells: cancer cells, marked in green, and T cells, highlighted in red. This dual-annotation approach not only aids in identification but also facilitates a deeper understanding of the interactions and responses under varying experimental conditions.

One of the primary goals of our study is to present an extensive collection of microwell images, with the total number exceeding 6000 groups. This rich dataset allows for an in-depth analysis of inherent characteristics across different groups. Such comprehensive data is crucial for accurately categorizing each group of images into appropriate clusters based on their intrinsic features.

A notable aspect of our dataset is the consistency in the size of the images. Each image within the dataset is uniformly sized, measuring precisely 173 *×* 173 pixels. This uniformity ensures that comparisons and analyses across different image groups are consistent and reliable.

The dataset’s depth and breadth, combined with the detailed annotations and uniform dimensions, make it an invaluable resource for our research. It enables us to trace and understand the nuances of cellular growth and behavior under various conditions, ultimately advancing our knowledge in this field.

### 4.3 Implement Details

We employed our methods using Pytorch and trained our model on a single V100-sxm2. During our experiments, we trained the model for 100 iterations, during training, the batch size was 16. The optimizer that we used is SGD. The initial learning rate was set to 1*e −* 4. The weight decay parameter was 1*e −* 4.

### 4.4 Evaluation Metrics

To comprehensively assess the outcomes of our feature extractor, we employ a diverse array of evaluation metrics, including accuracy(Acc), sensitivity(SE), specificity(SP), precision(PC), jaccard similarity(JS), and dice coefficient(DC).

### 4.5 Biological Interpretations

#### 4.5.1 Rationale for BCMA-targeting CAR Library Design

To accurately assess the capacity of the Enrich Trovo to identify highly functional CAR T cells from a pool of candidates, it was important to screen a library of CAR variants that all share identical target specificities but differ only in their proliferation and cytotoxic function. To enable this, our team designed a library of four candidates, schematized in Fig. 4 and equally represented within the library, that all leverage a single, previously identified and clinically validated scFv binder targeting B cell Maturation Antigen (BCMA), a B-cell specific antigen expressed preferably on mature B lymphocytes that serves as a promising and clinically relevant target in multiple myeloma [29]. While the same scFv, targeting BCMA, confers identical target specificity to all four CAR effector cells within the library, each library member differs in the features that comprise its intracellular signaling domain and its resultant ability to activate and initiate downstream signaling. To generate a broad range of T cell activation profiles, our library was composed of both non-signaling as well as first- and second-generation CAR designs. CAR B, one of the four CAR candidates in the library is a first-generation CAR design that includes only a CD3z signaling domain. However, a single antigen-dependent signal is usually inadequate for complete T cell activation, and T cells become anergic and do not proliferate in response to the antigen if they receive only this signal[30]. Consequently, given the single activation signal encountered by first generation CAR T cells, these CAR effectors are well known to have limited signaling that restricts their ability to induce durable T cell responses or prolonged cytokine release, ultimately attenuating their anti-tumor activity and target-induced proliferation [31][32]. The coupling of additional costimulatory signaling domains, namely 4-1BB or CD28, as with CAR A and CAR C, are known both preclinically and clinically to enhance the activation, survival, and effective expansion of CAR T cells [31][33][34]. In particular, Lisocabtagene maraleucel, a clinically validated CAR T cell therapy for patients with relapsed or refractory multiple myeloma, utilize a same design as CAR C [35]. Despite the impressive antitumor activity of second-generation CARs containing both 4-1BB and CD28 costimulatory domains observed in numerous preclinical studies, there are limited mechanistic studies that directly compare the performance of CD28 and 4-1BB CAR effector cells in parallel [36]. Broadly, though not exclusively, these studies seem to suggest that T cells that express CARs containing a CD28 costimulatory domain generally produce larger amounts of cytokines, including IFNy, IL-2 and/or TNFa, than CAR T cells costimulated with 4-1BB, suggesting some differences in initial proliferation and cytotoxicity [36][37][38][39]. Neverthless, greater persistence of CAR T cells with 4-1BB costimulation has been observed in some but not all mouse models [36][39]. However, in all these studies, a clear superiority of 4-1BB or CD28 costimulation has not been conclusively demonstrated. Furthermore, the ability to draw comparisons in these studies has been limited by variability not just in the costimulatory domains between CAR candidates but also in other structural elements of the CAR, such as the hinge and transmembrane regions, that can affect both cytotoxicity, proliferation and persistence [40]. In our library, both second generation CARs, CAR A and CAR C, were designed with identical binders as well as identical CD28 transmembrane domains to ensure that they only differed in the costimulatory signal. As a final control, our library also included a nonsignaling CAR effector cell, labeled ‘del’. These CAR effectors have intact extracellular domains but lack any intracellular signaling domains, as shown in Fig.4 and were not expected to show either cytotoxic activity or substantial proliferation. We consistently observed rapid initial reduction del T cells and significant cancer cell proliferation, damaging microwells during imaging. Thus, this sample was not included in the AI analysis.

**Figure 4:**
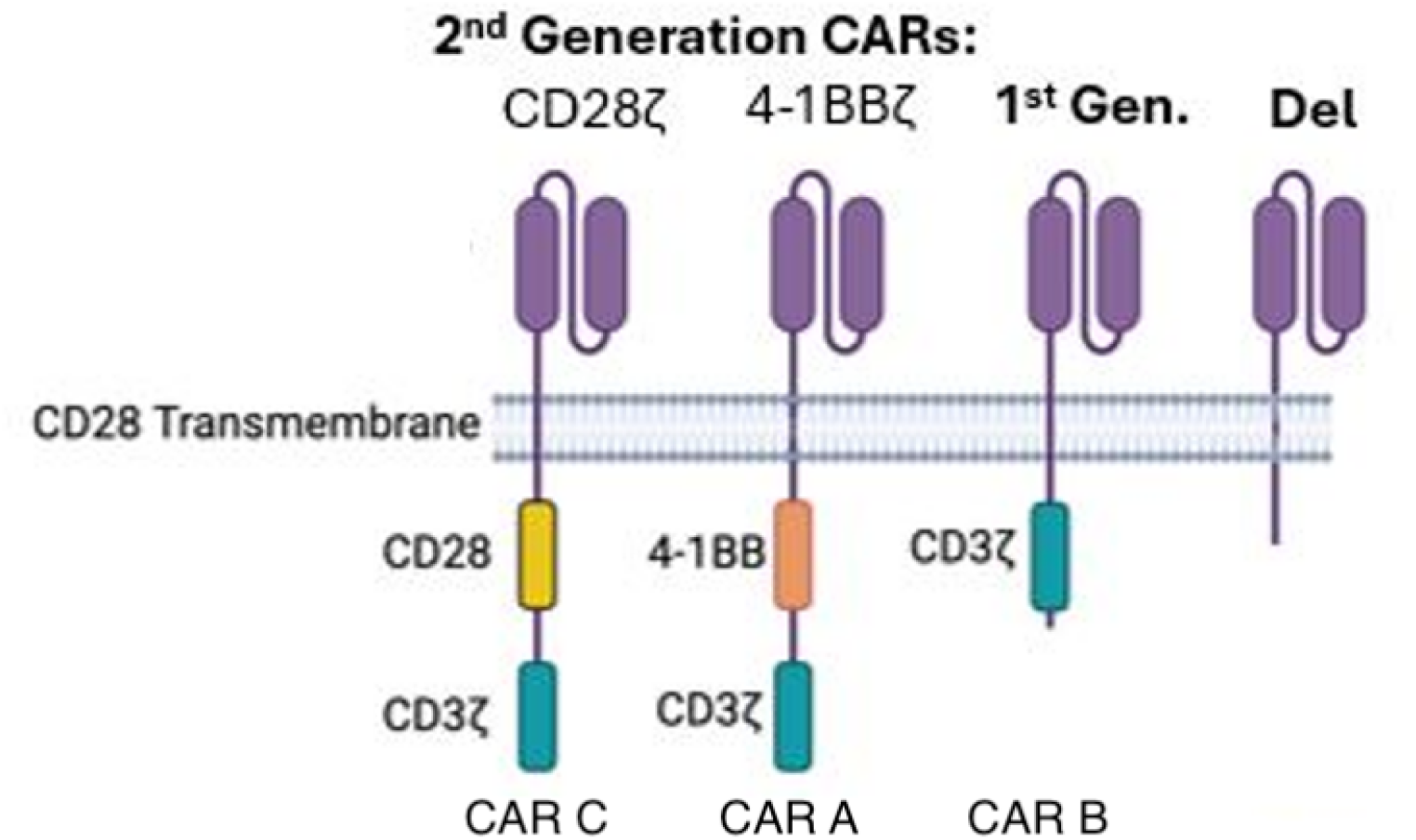
A schematic of the four-member CAR library utilizied in semi-pooled screening on the Trovo and subsequent image analysis. Each of the four library members contains a BCMA targeting scFv, that is connected to a CD28 transmembrane and intracellular signaling elements via a spacer sequence. The two second generation CARs, CAR A & B, contain a costimulatory domain (CD28 & 4-1BB respectively) in addition to CD3z. The first generation CAR, CAR C, contains only the CD3z and CAR D, a del CAR, lacks functional intracellular signaling domains.

**Figure 5:**
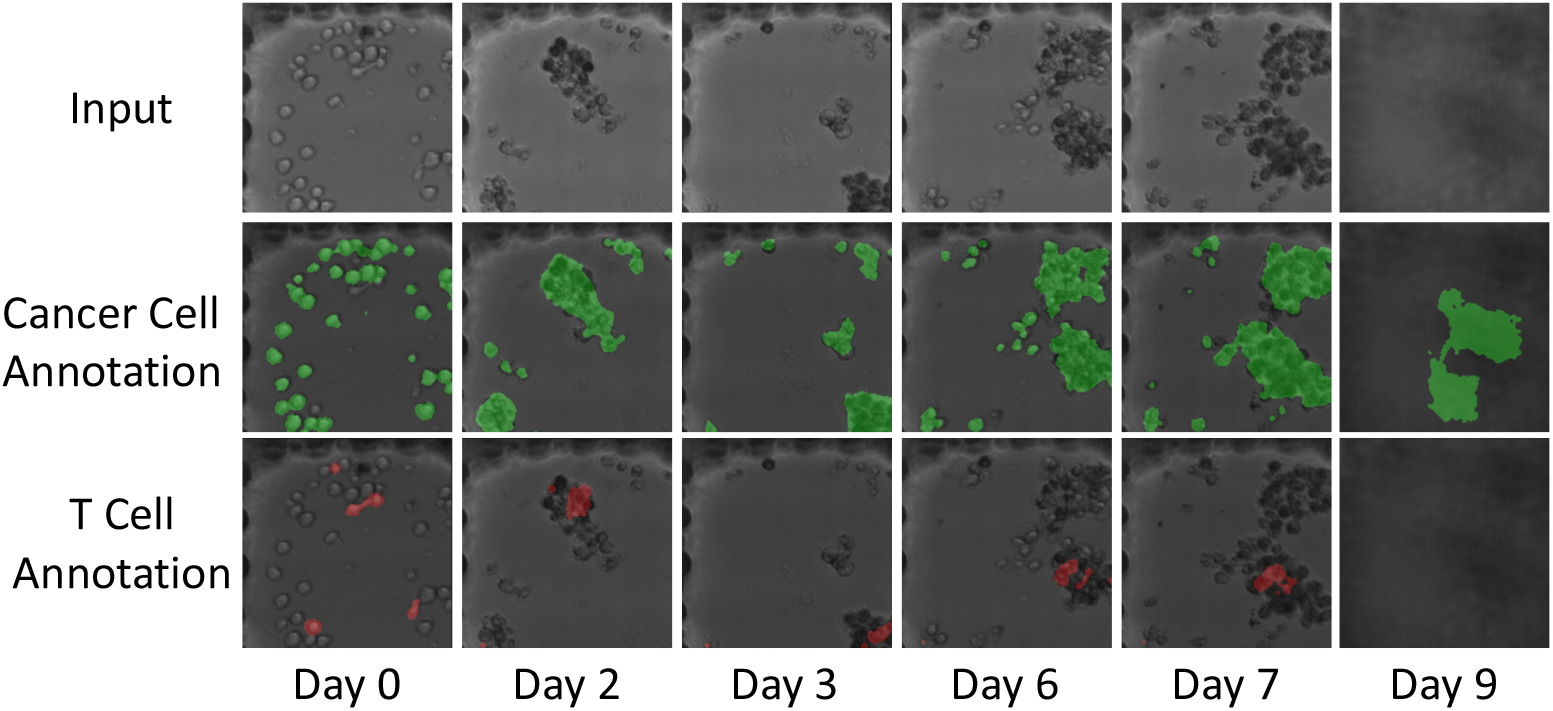
Overview of our dataset. Our dataset contains over 6000 groups of images. Every group consists of 6 images collected on different days. For every input image, we provide two different annotations: one is the cancer cells, and the other is the T cells.

#### 4.5.2 Correlation between unique groups found by AI

Being able to use high-throughput screening approaches to concretely identify CAR T clones with desired proliferative or cytotoxic phenotypes is crucial in shortlisting constructs to move forward for future studies. In this study, 4,000 “movies” were analyzed using the described AI algorithm from each of the following data sets: CAR A, CAR B, CAR C, and Pooled. Based on criteria set within the AI algorithm, 28 groups were formed based upon similar trends identified using the “movies” and the percent distribution for each construct was graphed (Fig.6 A). These groupings helped to identify unique differences between the CAR T cell clones aiding in identifying which microwells to capture. For example, in groups 8 and 27, CAR A is shown to be the most prominent. The derived trend lines (CAR T in red and cancer cells in green) based upon the “movies” in these two groups show that the CAR T cells grow very little but suppress cancer cell growth (Fig.6B). In contrast to this phenotype, CAR B, which is most prominent in group 18 has no CAR T cell proliferation and increased cancer cell proliferation based upon its trend lines (Fig.6 C). This might suggest that this CAR T construct cannot effectively prevent cancer cell outgrowth. Furthermore, upon looking at groups 6, 11, 21, and 22, where CAR C is most prominent, it can be observed that three of the four groups have increased T cell proliferation, and slow cancer cell outgrowth. However, unlike with CAR A there is not a decrease in cancer cell outgrowth. It can also be observed that in group 11, where CAR C is most prominent both the number of CAR T cells and cancer cells drop dramatically (Fig.6 D). This is likely due to the microwell starting with more CAR T cells comparatively, overwhelming the cancer cells and effectively killing them. Without enough cancer cells to stimulate CAR C these cells were unable to continue to proliferate. In addition to identifying unique groupings for each of the constructs, the algorithm also identified groupings where two of the three or all three of the constructs were distributed relatively evenly (Fig.6A). This can be seen in groups 3 and 4 where both CAR A and CAR B are most prominent, and similarly in groups 26 and 28 where CAR A and CAR C are most prominent (Fig.6A). In many of the assigned groups all four of the data sets are equally observed or there are not enough “movies” binned into that specific group to confidently determine a unique phenotype. In turn those groups would not be selected for further processing of the CAR T cells. To identify the cells to capture, CAR T cells are plotted against cancer cells on a plot differentiation graph. In this example the average cell confluency for each group is plotted as shown in Fig.6 E and Fig.6 F. On Day 1 there is no noticeable trend in the clustering areas for each of the CARs. However, by day 10 each of the groups where there is a prominent CAR have clustered together except for CAR B. The groups with a significant representation of CAR A clustered together on the graph and showed suppression of cancer cell growth but slow CAR T cell proliferation, similar to what was seen in the trend lines of the individual movies. Similarly, the groups that were prominent for CAR C also clustered together showing suppression of cancer cell growth and increased CAR T cell growth. Increased CAR T cell growth - a feature that is supported by the trend lines in the representive movies for each group. CAR B didn’t cluster very well, and this is likely because this CAR is overall not being stimulated by the cancer cells and dies or persists in the microwell but does not suppress cancer cell growth. The lower left region of Fig. 6 shows two groups (orange and green) both are good killers. Orange/CAR A shows less T cell proliferation and green/CAR C shows high T cell proliferation, which is consistent with the CAR T cell design.

**Figure 6:**
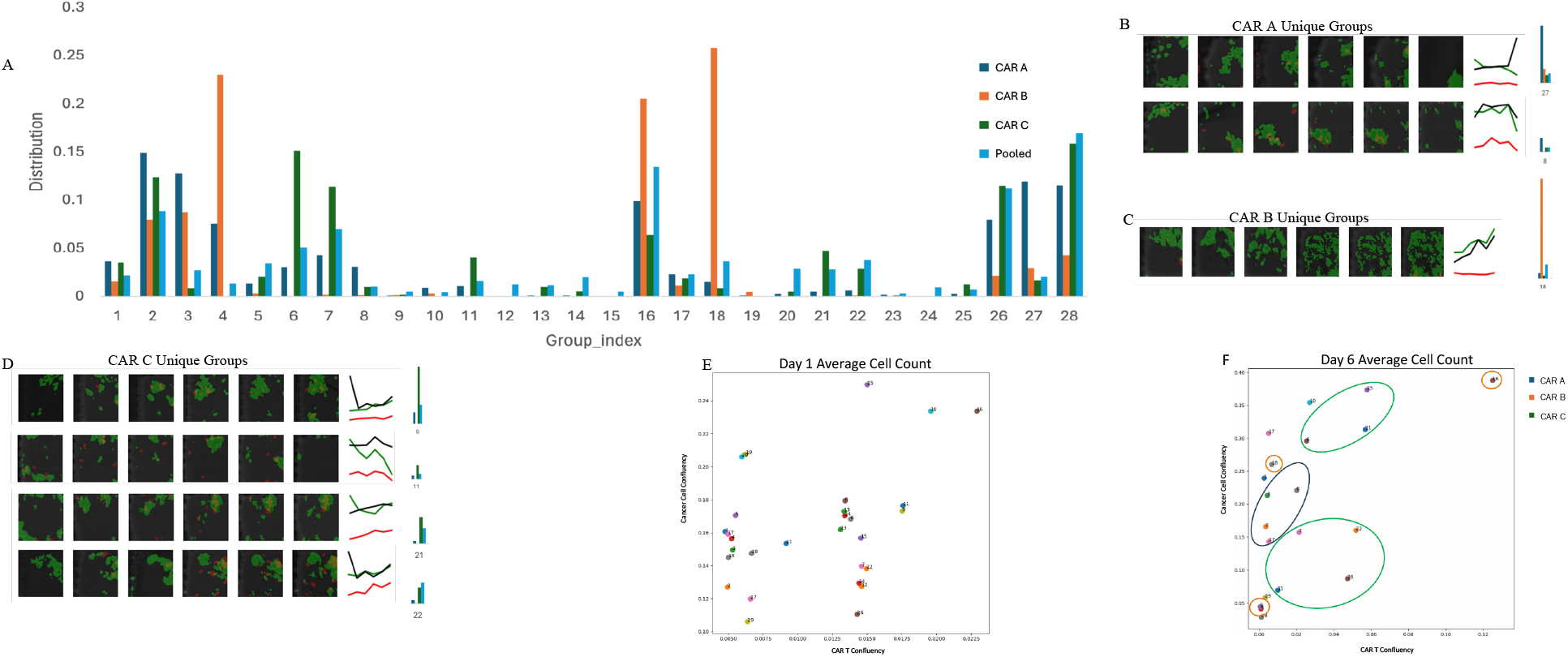
Using the AI algorithm groups, 16,000 “movies” were analyzed and distributed based on similar trends. (A) Criteria set by the AI algorithm formed 28 unique groups using data from the following data sets: CAR A, CAR B, CAR C, and Pooled. (B, C, D) A representative “movie” for each of the groups where the CAR T cell lines CAR A, CAR B, and CAR C are most prominent. (E, F) For each of the 28 unique groups the average confluency of cancer cells and CAR T cells were determined and plotted against each other on day 1 or day 6. On day 6 clusters for each of the CARs are identified by a representative circle.

## 5 Comparison with other models

Given the relative novelty of this subject, a direct comparison between our approach and existing methods is not readily feasible. To provide tangible quantitative insights underscoring the effectiveness of our innovative architecture, we undertake a systematic approach. This involves adapting several alternative architectures to align with our model’s framework, thereby enabling a comparative analysis.

The tabulated outcomes, as shown in Table 1, we provide the results obtained from alternative methods like Resnet[27], HRNet[28], U-net[7], alongside those yielded by our proposed approach. Upon examination, it becomes evident that our proposed methodologies outperform their counterparts, consistently demonstrating superior performance. This comparative evaluation substantiates our claim of attaining the most effective outcomes within the context of the presented problem domain.

**Table 1:** Quantitative outcomes of various architectures, with the top results emphasized in **bold**.

| Methods | Acc | SE | SP | PC | JS | DC |
| --- | --- | --- | --- | --- | --- | --- |
| VGG ([26]) | 92.64 | 84.79 | 95.68 | 93.47 | 85.26 | 90.47 |
| Resnet18 ([27]) | 93.52 | 86.73 | 96.54 | 94.27 | 87.24 | 91.82 |
| Resnet34 ([27]) | 94.05 | 87.26 | 97.15 | 96.18 | 88.82 | 92.63 |
| HRNet ([28]) | 95.39 | 89.47 | 97.18 | 97.49 | 88.67 | 94.18 |
| U-net ([7]) | 94.25 | 88.15 | 97.02 | 95.14 | 89.21 | 92.10 |
| Ours | **97.84** | **91.45** | **99.03** | **98.87** | **90.56** | **95.06** |

## 6 Ablation Study

### 6.1 Effectiveness of the preprocessing module and temporal features

In order to showcase the effectiveness of both our preprocessing module and the integration of temporal features, we did ablation studies to demonstrate this behavior. As shown in Table 2, it becomes evident that introducing preprocessing and temporal features yields a discernible enhancement in our model’s performance. This augmentation is attributed to the distinct advantages offered by these two components.

**Table 2:** Ablation study for the existence of preprocessing and temporal features, with the top results emphasized in **bold**.

| Methods |  | Acc | SE | SP | PC | JS | DC |
| --- | --- | --- | --- | --- | --- | --- | --- |
| Preprocessing | Temporal Feature |  |  |  |  |  |  |
| × | × | 94.37 | 88.14 | 95.47 | 94.58 | 88.32 | 92.15 |
| ✓ | × | 95.46 | 89.57 | 96.37 | 95.47 | 88.93 | 93.16 |
| × | ✓ | 96.36 | 90.47 | 97.47 | 97.24 | 89.48 | 94.19 |
| ✓ | ✓ | **97.47** | **91.84** | **98.14** | **98.26** | **90.94** | **95.59** |

The preprocessing phase is particularly effective in enabling our model to refine its focus on foreground elements, allowing a more accurate analysis of relevant aspects. In parallel, the incorporation of temporal features amplifies the informational richness compared to individual images. The comprehensive analysis provided by these temporal features significantly contributes to the model’s performance, as corroborated by the results presented.

### 6.2 Effectiveness of human-designed features

Human-designed features are important in our feature extractor. Within this section, we conducted an ablation study to substantiate the efficacy of our human-designed features. The results, as highlighted in Table 3, illustrate that incorporating human-designed features yields a discernible improvement in our model’s performance. Delving deeper, the rationale behind this enhancement lies in the unique attributes of these human-designed features. Their ability to contribute statistical features that are intricate and not readily learnable by neural networks imparts a decisive advantage, thereby bolstering our model’s overall performance.

**Table 3:**
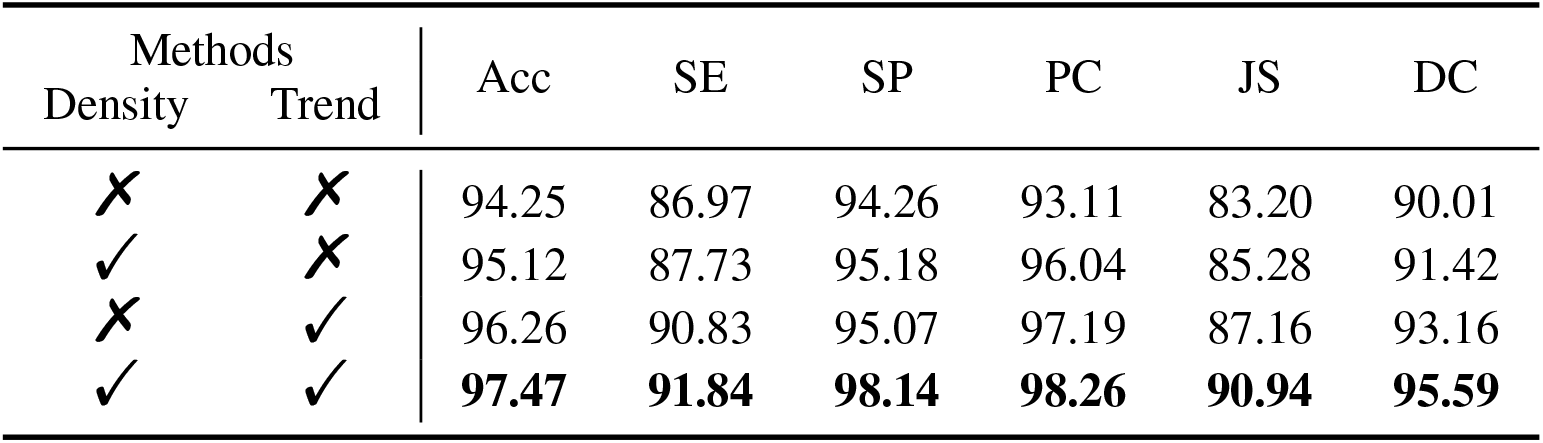
Ablation study for the existence of human-designed features, with the top results emphasized in **bold**.

| Methods |  | Acc | SE | SP | PC | JS | DC |
| --- | --- | --- | --- | --- | --- | --- | --- |
| Density | Trend |  |  |  |  |  |  |
| × | × | 94.25 | 86.97 | 94.26 | 93.11 | 83.20 | 90.01 |
| ✓ | × | 95.12 | 87.73 | 95.18 | 96.04 | 85.28 | 91.42 |
| × | ✓ | 96.26 | 90.83 | 95.07 | 97.19 | 87.16 | 93.16 |
| ✓ | ✓ | **97.47** | **91.84** | **98.14** | **98.26** | **90.94** | **95.59** |

## 7 Results of our explanation module

As seen in Fig.7, the model highlights the region of the image where the model focuses. Based on the fact that the highlighted region matches the cells, it suggests that our model attempts to capture cell-specific information. As shown in Fig.8, our methods can not only explain each cluster but also provide a comparison between a given cluster and the others.

**Figure 7:**
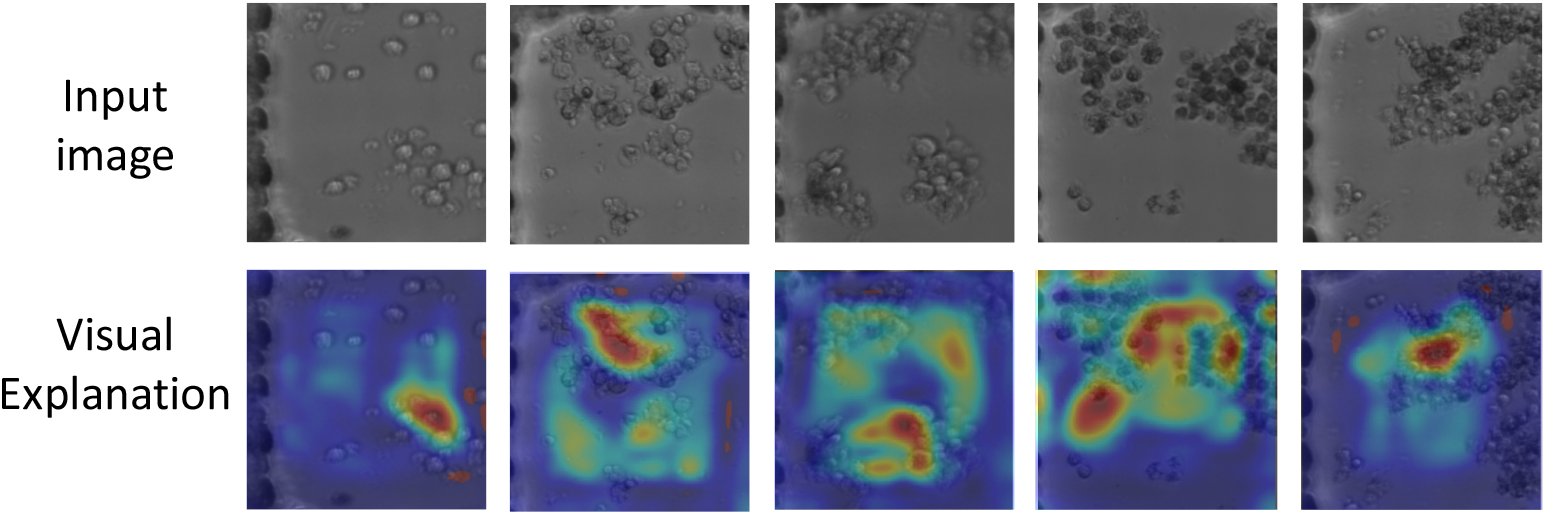
Our methods can highlight the region where the model focuses on and visualize them using an attention map

**Figure 8:**
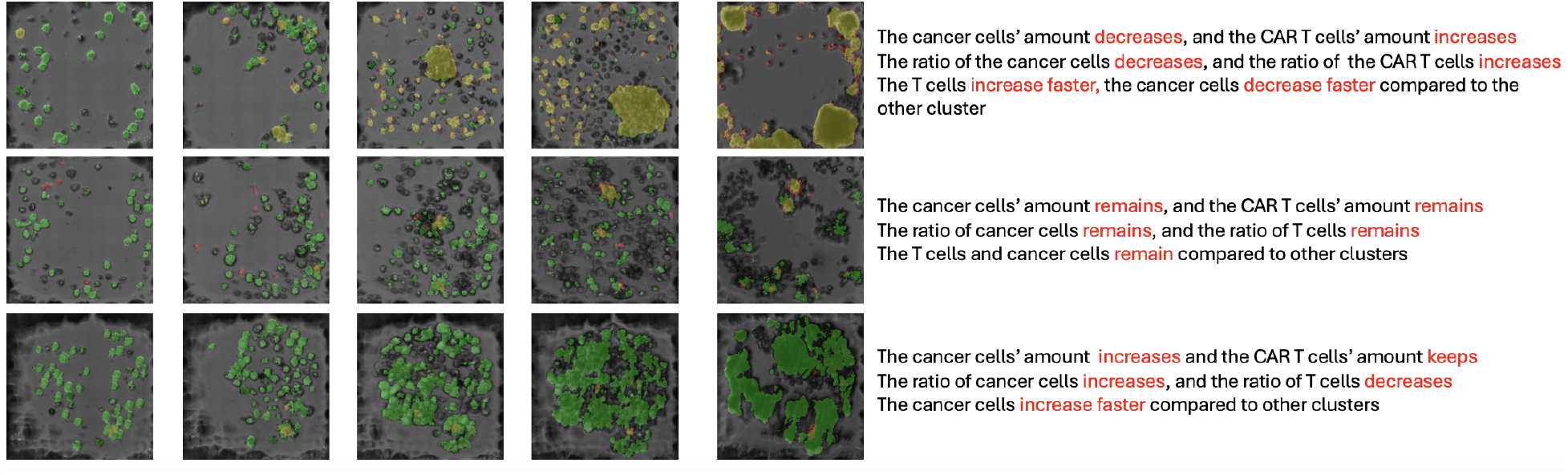
Our methods can provide the explanation for each cluster and a comparison between this cluster and other clusters. Here, we randomly pick one group of images for every cluster.

## 8 Intra and Inter covariance matrix

In order to provide a more comprehensive illustration of the validity of our cluster algorithm, we present the intra and inter-covariance matrices in this section, as depicted in Fig.9. The intent behind showcasing these matrices is to elucidate the effectiveness of our clustering methodology through visual representation.

**Figure 9:**
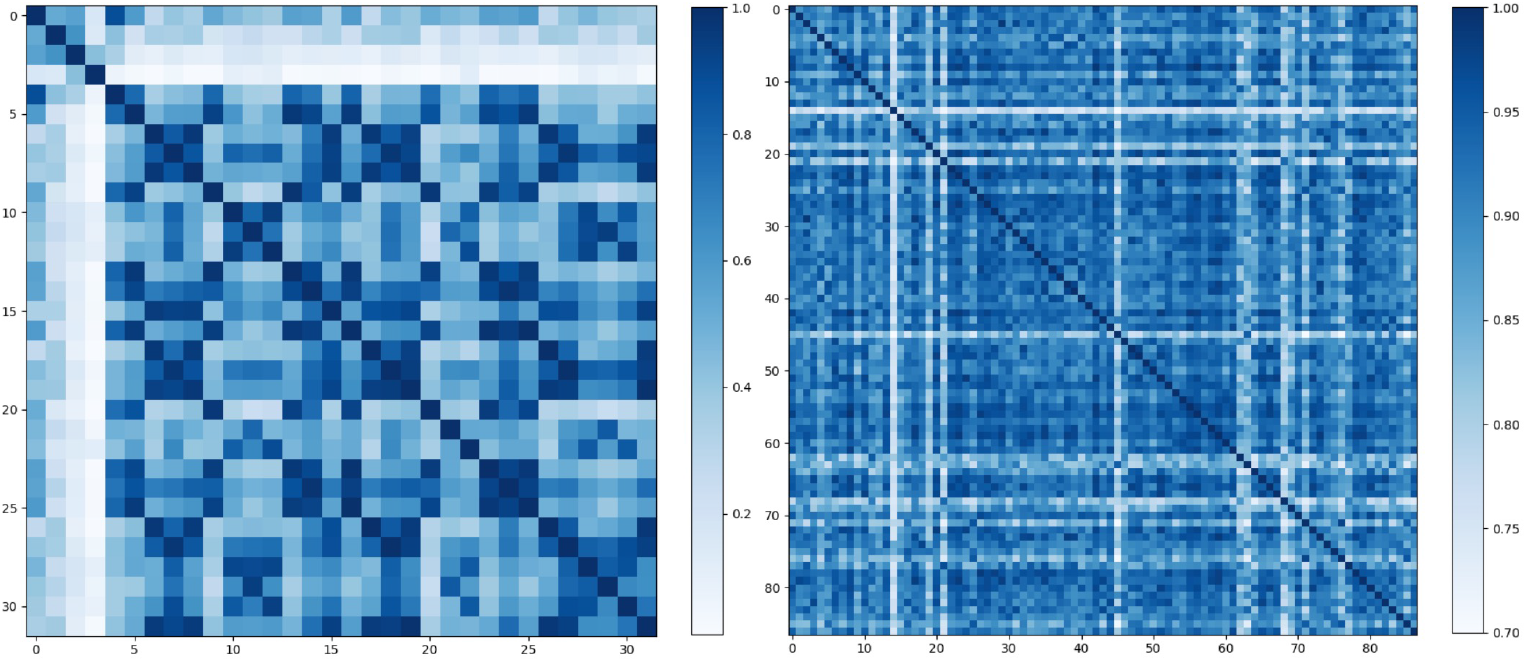
We visualize intra and inter-covariance matrices for different clusters. The left image is the inter-covariance matrix, with low similarity between different clusters. The right image is the intra-covariance for a random cluster. Notice that the start point for the right image is 0.7, meaning that every image in this cluster has high similarity.

For the calculation of the intra-covariance matrix, we adopt a methodical approach. Commencing with the random selection of a cluster, we proceed to compute the cosine similarity between the feature vectors of every pair of images within that cluster. The outcomes are then presented in the form of a matrix, effectively encapsulating the cosine similarity between various image pairs. A notable trend emerges, as observed in the bottom image of Fig 9: images within the same cluster exhibit significantly higher cosine similarity, a testament to the accuracy of our clustering approach.

The process for generating the inter-covariance matrix follows a similar methodology. The starting point involves the calculation of cluster centers for each individual cluster. Subsequently, the cosine similarity between these cluster centers is determined. As depicted in the left image of Figure 9, different clusters manifest relatively low cosine similarity, reaffirming the distinctiveness of our clustering outcomes.

The visual representation of these covariance matrices provides a compelling visualization of the efficacy of our cluster algorithm. Inter-cluster similarity remains low after clustering, while intra-cluster similarity remains notably high. This observation underscores the robustness of our clustering approach in discerning distinctive patterns and grouping images effectively.

## 9 Conclusion

In summary, we have introduced an innovative approach that acquires information from individual images and captures cellular dynamics. Following the extraction of features, our method employs a clustering algorithm to categorize each image group into distinct clusters. The absence of the labeled data poses a challenge in evaluating our clustering algorithm’s performance quality.

To address this, we have integrated two distinct explanation modules. By leveraging textual and visual explanations, we have developed an understanding of the underlying reasoning behind the model’s predictions. This multifaceted approach enables us to obtain valuable insights into the effect of different CARs.

We believe that the versatility of our proposed methodology extends beyond the confines of the clustering of images of microwell cocultures for hit selection. With its potential to help researchers better understand the influence of different environments on cancer cells and T cells, we anticipate that our approach will find applications in diverse areas within drug development while delivering quality performance.

